# Paternal grandparental exposure to crop failure or surfeit during a childhood slow growth period: Epigenetic marks on grandchildren’s growth, glucoregulatory and stress genes

**DOI:** 10.1101/215467

**Authors:** Lars Olov Bygren, Patrick Müller, David Brodin, Gunnar Kaati, Jan-Åke Gustafsson, John G. Kral

## Abstract

This latest in our series of papers describes transgenerational methylation related to midchildhood food availability in 19^th^ century Överkalix, Sweden. Failed vs. bountiful crops differentially influenced methylation in grandchildren of paternal grandparents exposed to feast or famine during their Slow Growth Period (SGP), a sensitive period preceding the pre-pubertal growth spurt. In this case-study of 8 tracked 75-year old progeny with differential ancestral exposure, we found, in 40 posited gene ontology pathways, 39 differentially methylated CpG regions (DMRs) related to famine, excess food and food-insecurity stress, 9 of which with DMRs above 5%. Three gene ontology terms (GOs) “insulin processing”, “adipose development” and “hypothalamus development” were key, with DMRs >14%. An unbiased test of known pathways revealed four nuclear transcription factors upstream of promotors repressing the pathway following paternal grandparental famine experience, as well as 4 upregulated GOs with average DMRs >20%. We conclude that this is the first demonstration of human transgenerational inheritance of epigenetic marks following ancestral childhood exposure to variable food availability inducing early developmental origins of adult disease.

---

Paternal grandparent food supply, preceding their prepubertal growth spurt, induced epigenetic marks in three gene pathways reflecting famine, excess food and food-insecurity stress. Our epidemiological findings of adverse transgenerational effects of ancestral overnutrition and conversely, beneficial effects of famine during SGP^1–6^ prompted us to study individuals whose paternal grandparents were randomly exposed during their Slow Growth Period (SGP) - a sensitive period preceding the pre-pubertal growth spurt - to failed or bumper crops potentially engendering epigenetic marks on mechanistic molecular gene pathways.

For detailed analyses of specific gene ontology (GO) pathways, we tested three posited structural classes of genes associated with each of 40 preselected GO terms (http://amigo.geneontoloev.org) for differentially methylated CpG regions (DMRs) between the experimental groups. We selected GO pathways with average methylation differing >5%. The posited pathways comprised glucoregulatory genes interpreted as immediate sensors of decreased glycogen stores, whereas those of lipids and ketones were posited to reflect the duration and magnitude of the famine or alternatively, responses to overnutrition. Food insecurity stress was assumed to have activated the HPA-axis. We focused on two putative response lines in direct descendants from a) grandfather to son with grandson and b) paternal grandmother to son with granddaughter, adjusted for crop size during famine or surfeit. We found 39 DMRs in grandchildren after grandparental differential exposure to food availability in GO-pathways associated with famine, excess food and environmental stressors. Nine of them were large, three GOs exhibiting remarkable DMRs in grandchildren (Table 1). An unbiased analysis detected transcriptional pathways, such as CCAAT-box (AKA C/EBPbeta) affecting adipogenesis, laying down fat, the primal tissue for energy storage.

**Table 1.**
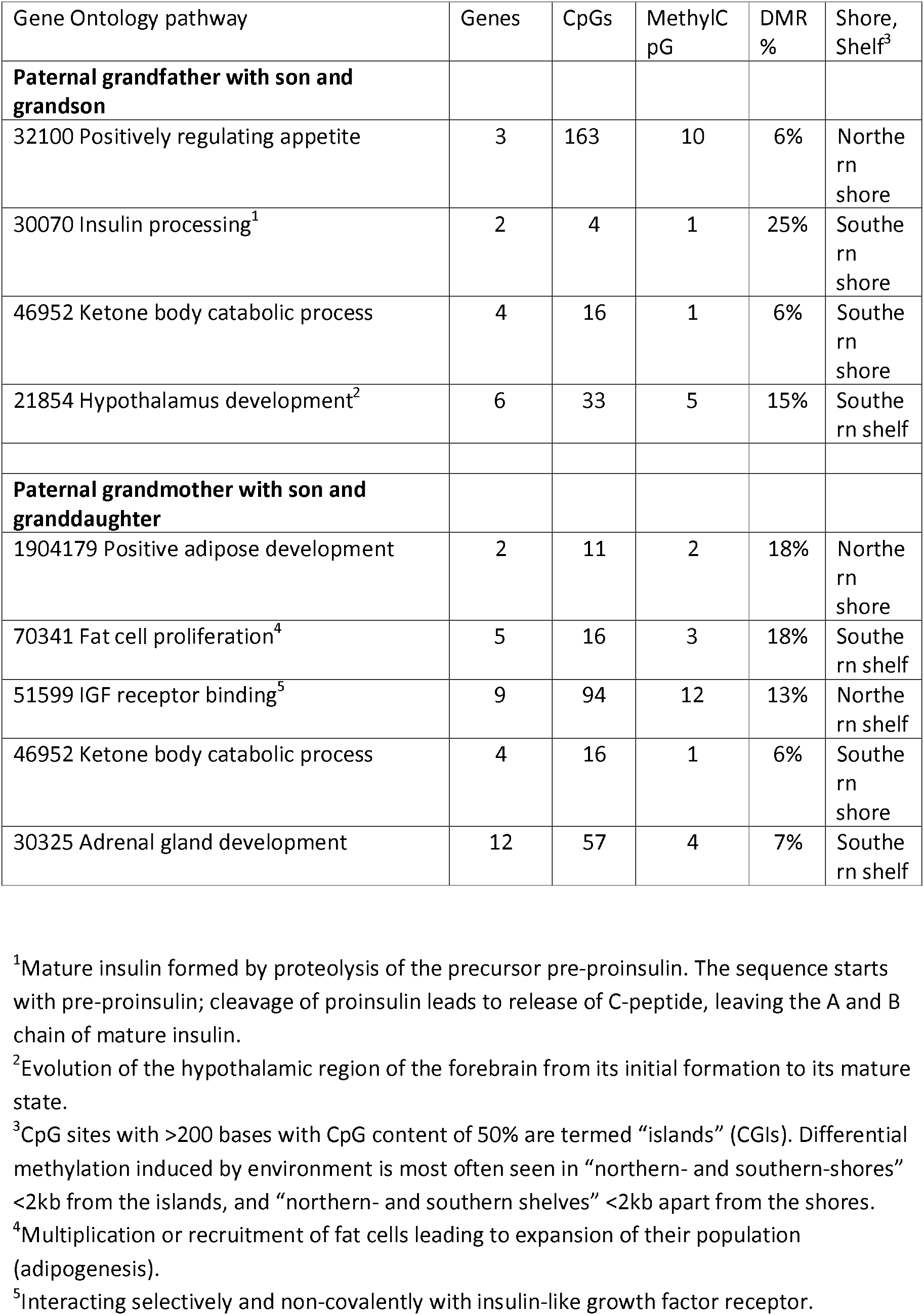
Gene ontology pathways related to hunger, excess food and stress associated with paternal grandparental exposure to variable food availability with DMRs in grandchildren.

## RESULTS

Exploiting an “experiment” of nature, we studied DNA methylation in grandsons and granddaughters in the paternal lineage. We found 39 DMRs among 40 posited pathways in *islands*, North and South *shores* and North and South *shelves* related to famine, nutrient excess and stress, among which 9 pathways had DMRs above 5%, and 3 above 14%. The other 31 pathways had 0-5% DMRs. An unbiased test of pathways in *islands* revealed 2 pathways with DMRs 40% and 2 with DMRs 22 and 21% (Table lb).

**Table 1b.**
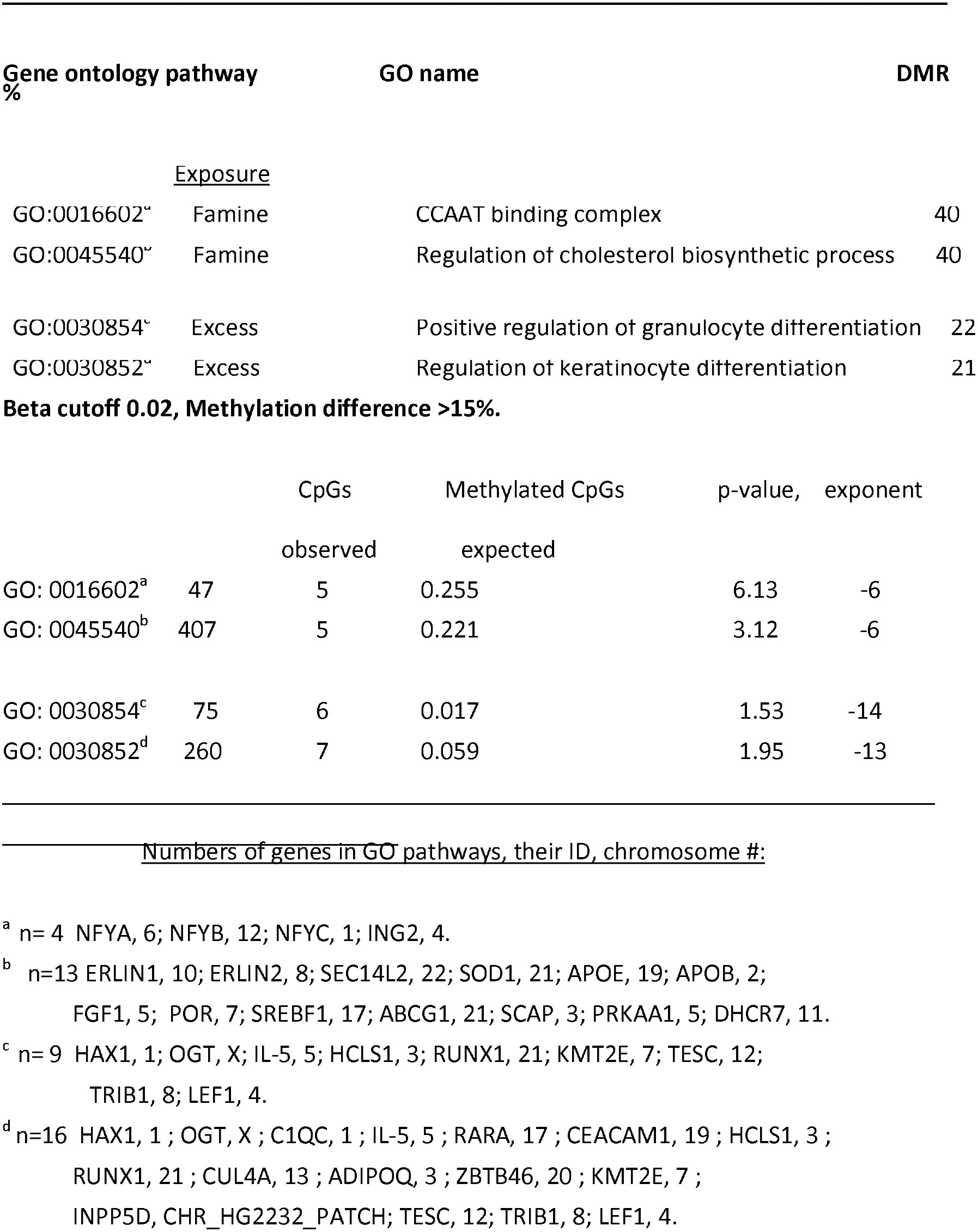
Gene ontology (GO) pathways in 8 grandchildren (4 men, 4 women) whose paternal grandparents were exposed to famine or food excess.

### Paternal grandpa rental exposure vs. methylated genes in adult grandchildren

The main metabolic pathways implicated in grandpaternal SGP exposure to food-insecurity stress transferred to grandsons were “appetite”, “insulin processing”, “ketone body catabolic process” and “hypothalamic development”. Differential grandmaternal exposure to feast or famine was associated with marks in “adipogenesis”, “insulin-like growth factor receptor (IGF-R) binding”, “ketone body catabolic process” and “adrenal gland development” (Table 1). The probands’ 4 ketone body catabolism genes (GO 30070) were the same for grandfathers’ as for grandmothers’ exposures, supporting the findings. The directions of the differences were intuitive: famine decreased energy-demanding pathways whereas overnutrition enhanced them. Altogether three pathways were key.

#### Three key gene ontology terms implying transgenerational epigenetic inheritance

<u>GO 0030070: “insulin processing”</u> modulates insulin production downstream from proteolysis of the precursor preproinsulin via C-peptide. Proinsulin is cleaved to release C peptide. This pathway had a DMR in the male line in the Southern Shore involving 2 genes: PCSK2, proprotein convertase (also called NE2 HUMAN) in chromosome 20 and CPE Carboxypeptidase E (also called CPBE HUMAN) in chromosome 2. This pathway is related to GO 1901142: “insulin metabolic process”.

<u>GO 1904179: “adipose development”</u> includes gene NR1H4 (also called BAR;FXR;HRR1;RIP14) and the nuclear hormone receptor pthr 24082, “bile acid receptor”. Other genes are SIRT1 (also called SIR2L1) and chromatin regulatory protein sir2, pthr 11085.

<u>GO 0021854: “hypothalamus development”</u> involves GSX1 (GShomeobox1), PTX2 (Pituitary homeobox2), UBB (Polyubiquitin-B) and CRH (Corticoliberin; Corticotropin-releasing hormone).

#### Unbiased correlations (Table 1b)

Regardless of sex, the following gene ontology terms were highly correlated with famine:

<u>GO 0016602</u>: A transcription complex that binds to the CCAAT-box upstream of promotors typically consisting of three subunits, i.e. NFYa-c. Four genes were repressed: NFYAon Chromosome 6, NFYB on chromosome 12, NFYC on chromosome 1 and ING2 on chromosome 4.

<u>GO 0045540</u>: “Regulation of cholesterol biosynthesis” (formation of cholesterol) was repressed in 13 genes, such as ERLINI, chromosome 8 and 10; SECI4L2, chromosome 22 and SODI, chromosome 21.

Several cell differentiation pathways correlated with food surplus, notably the following two:

<u>GO 0030854</u>: “Positive regulation of granulocyte differentiation” available for expression of its 9 genes, e.g. HAXI on chromosome 1, OGT on the X chromosome, ILS on chromosome 5 and HCSL1 on chromosome 3.

<u>GO 00 45616</u>: “Regulation of keratinocyte differentiation” with 16 genes, such as HAXI, OGT, ILS and C1QC on chromosome 1.

#### Parental age data and selection

Average paternal ages at birth of first child were 32, 31 and 27 years in the three generations with grandparents exposed to famine and 34, 27 and 27 years with grandparents exposed to feast. Seven index persons were alive at 74, the eighth having died at 63 years. For grandparents and parents exposed to famine during SGP, the average ages of death were 61 and 78 years respectively whereas after feast they were 87 and 71 years (Table 2). Sequential genetics findings contributing to differential methylation were not taken into account.

**Table 2.** Reproductive fitness following grandparental famine or feast in the Slow Growth Period

| Index person | Age at first child's birth |  |  | Number of children |  | Longevity |  |  |
| --- | --- | --- | --- | --- | --- | --- | --- | --- |
| Exposure | Grandparent | Father | Grandchild | Father | Grandchild | Grandparent | Father | Grandchild |
| **Paternal grandfathers' exposure** |  |  |  |  |  |  |  |  |
| Famine |  |  |  |  |  |  |  |  |
| Index person No 15 | 39 | 38 | NA | 4 | NA | 60 | 69 | 74+ |
| Index person No 53 | 27 | 31 | 27 | 7 | 1 | 49 | 98 | 74+ |
| Feast |  |  |  |  |  |  |  |  |
| Index person No 14 | 28 | 23 | 30 | 6 | 3 | 80 | 70 | 74+ |
| Index person No 23 | 33 | 26 | NA | 11 | NA | 94 | 83 | 63 |
| **Paternal grandmothers' exposure** |  |  |  |  |  |  |  |  |
| Famine |  |  |  |  |  |  |  |  |
| Index person No 59 | 39 | 38 | 34 | 3 | 2 | 80 | 83 | 74+ |
| Index person No 86 | 24 | 20 | 22 | 6 | 2 | 55 | 64 | 74+ |
| Feast |  |  |  |  |  |  |  |  |
| Index person No 3 | 36 | 29 | 23 | 10 | 4 | 86 | 65 | 74+ |
| Index person No 22 | 35 | 31 | 29 | 4 | 2 | 89 | 68 | 74+ |

NA. Not available

## DISCUSSION

We propose that known adaptations to energy deficiency or excess in cells, organs or organisms are expressed transgenerationally, appearing as epigenetic marks in humans. Famine might induce marks around promoters in at least two types of pathways: one general or nutrient-specific, related to *diminished* substrate stores of glycogen, lipid or protein, or *excess*, nutritoxic exposure (cf. eutrophication in plants), the other to environmental stresses of famine through activation of the hypothalamo-pituitary-adrenal (HPA) axis, as in food insecurity.

In earlier papers we described a unique cohort of descendants of grandparents exposed randomly to these weather-dependent variations in food availability. Data emanated from detailed archival records enabling correlations between demographic and crop yield statistics^1–6^. Our finding of SGP sensitivity findings have been replicated in Germany and Sweden ^7,8^. Our current paper adds mechanistic information based on blood sampling of seven 75-year-old grandchildren and one 63 years old, enabling determination of candidate epigenetic markers reflecting differential methylation in gene pathways posited to be affected by exposure of their grandparents to crop failure versus bountiful harvest during the grandparental childhood SGP preceding the prepubertal peak stature growth. Changes in DNA methylation levels of cytosine and guanine (CpGs) differentially methylated in one generation could explain transgenerational epigenetic inheritance in the third generation^9^.

A 2015 review concluded that evidence suggesting that acquired epigenetic marks pass to the next generation was limited; many others have noted the absence of any mechanism by which gene-regulatory information is transferred from somatic cells to germ cells in the study of transgenerational epigenetic inheritance (review: Nagy and Turecki^10^). Our epidemiological and epigenetic results demonstrate sex differences, e.g. in development of preproinsulin as has been seen in human pancreatic is lets^11^. The sex-bound epigenetic inheritance has been interpreted simply as epigenetic actions and disease manifestations being sex specific, as is the case with many conventional genetic variants^12^. The transcriptions are also not fully understood. Most confusion emanates from not knowing how exposed somatic cells can communicate their exposure to the germline which induces changes lasting for generations.

It is difficult to understand or accept epigenetic inheritance if there is a real rather than just a theoretical barrier between somatic and germline cells. One route to overcome this barrier might be the proposal that extracellular nano-vesicles, neighboring germ cells, shed by somatic cells contain the material required for transcription, able to bypass the barrier^13^. In the SGP the male primordial germ cells form active spermatozoa whereas human ovarian stem cells that can be modified in the SGP probably are required.

Presently the preponderance of evidence suggests transgenerational cumulative effects of exposures: diet, behavior, environmental chemicals, activity and the microbiome. For reviews, see Sales et al.^9^ and Vaiserman et al.^14^.

We have focused on the paternal line to discern transgenerational epigenetic inheritance disentangled from maternal-fetal and maternal-infant influences. The pathways in the paternal line are probably similar in the maternal line. We found three interesting gene pathways influenced by paternal grandparents’ exposure resulting in grandchildren’s DMRs, pathways induced by famine, overnutrition and food insecurity stress. When the paternal grandfather had been exposed to famine, the grandson exhibited DMRs of insulin processing. When the paternal grandmother had been exposed the granddaughter had DMRs of GOs related to environmental stress such as “hypothalamus developmenť(influencing the HPA-axis) related to increased female susceptibility to stress, yet explaining the benefit to mental health recently found in a different setting^7^.

A strength of our study is that it exploits a natural secular phenomenon viz. variable food security, the effects of which are documented in a homogeneous well-documented 19^th^ century population, allowing exceptional tracking of ancestors. Few founders colonized the area 500 years earlier whereby the area metaphorically became an island of native speakers of Swedish surrounded by Sami- and Finnish-speaking neighbors. Furthermore the index persons, the grandchildren, are dispersed all over Sweden^5^ and have neither been exposed to famine nor, being born before WW2, experienced current food excesses during their childhood, preceding the current obesity epidemic. Differences in blood tissue composition or heterogeneity of blood probably were of minor importance in this very homogenous population.

A second strength is the focus on two lines of inheritance that emerged in the epidemiological studies, the line from the paternal grandmother’s son and his daughter and the line from the paternal grandfather with son and grandson. Thirdly our *a priori* selection of gene ontologies, GOs, based on our earlier epidemiological findings of cardiovascular morbidity related to food security reduces the risk of spurious correlations. A confound of multiple testing could be suspected having 40 GOs but the ontologies were chosen for their evidence-based responses to only three potential pathways: feast, famine and stress. In addition, our unbiased test identified related transcriptional genes.

Our earlier published studies in the community and correlation of age at the birth of the first child, number of children and variance of survival over three generations ruled out that variable availability of food during SGP was caused selectively^2^. In human studies, measures are very crude but the figures for the eight subjects do not demonstrate any biased selection owing to ancestors’ random exposure during the SGP. The main weakness is having only 8 index cases, with low statistical power. Furthermore, seven were survivors up to age 75 years (one to 63) during which their own exposures might have induced methylations confounding those inherited from the grandparents and parents. On the other hand, epimutations analogous to mutation most often do not appear on the clinical horizon until late in life. The subjects were sufficiently healthy to participate and travel to the nearest health facility to give blood.

Confounding could have occurred through differences in exposure to the feast and famine between index cases owing to social circumstances such as belonging to a family with better food resources, or to other mechanisms differentially affecting allele-specific or other variables’ methylation^12^. However, the natural experiment exposed all families with little variation in poverty in the 19^th^ century mitigating this confound, as supported by our earlier research in the community^4^. The isolation of this community during the centuries up to the present might have resulted in a relatively inbred cohort and diminished assortative mating, both benefitting the epigenetic analysis. We present a case series of epigenomes of 8 adult grandchildren with archival data exhibiting their paternal grand-paternal or -maternal exposure to crop failure or large crops during their SGP linked to differential methylation of genes associated with energy balance and hypothalamic development. The salient novel finding is that grandparental childhood exposure was reflected in grandchild profiles of CpG methylation of key genes. Our epidemiological data from the cohort from which these strategic cases were drawn strengthens the molecular findings and indicates possible mechanisms of inheritance that might be important for the field. The presence of epigenetic changes attributable to environmental influences potentially engendering several prevalent multifactorial chronic diseases and conditions. These findings replicated in other settings may provide guidance for preventive and therapeutic interventions targeting methylation.

## METHODS

### Classification of exposure to food availability in the environment

A seven point scale for estimating annual crop sizes was employed during 1816-1849 [Statistics Sweden (Tabellverket)]^15^. It spanned from “total failure” to “abundant” crops and has been used since in demographic research. For the years 1865-1902, birth years of the 8 grandparents, statistical tables of crop yields were the primary source, validated against food price statistics^16^ and qualitative reports from a gubernatorial *Crops and National Aid Office* (Hushållningssällskapet). Using these sources, we created the ordinal seven-item “Hellstenius scale”. Over the years the criteria for the items of the scale have changed apace with changes in the Swedish language. The current translation is as follows: Total failure (0), Sparse general growth (1), Weak or Small (2), Below average (3), Average or Mediocre (4), Above average (5) and Good or Abundant (6). We defined 0-2 as famine, 3-4 as mediocre, and 5-6 as excess food availability. We used May 1 of the year following crop failure as the time of least food availability in this area around the 66^th^ latitude and determined November 1 of the year following good harvest with pig and cattle slaughter when the meat could be frozen, as the time of greatest food availability (excess,eutrophic). Grandparent age on those dates was used to relate food availability to his or her SGP.

### SGP, the slow growth period

Two developmental periods in our first analysis were posited to differ in sensitivity to food supply affecting transgenerational responses: the prepubertal growth peak and the preceding slow growth period (SGP). The growth peak was seen as a sensitive period for a child to be exposed to famine. The slowest growth period before the peak was posited to be less prone to energy variation. These two periods were set in relation to puberty, taking into account individual variation, nutrient availability shifting the timeframes, and realizing that the age distribution of puberty was skewed to the right during the 19^th^ century compared to later. The two periods were derived by superimposing the stature growth velocity curves of Prader et al.^17^ upon the ages of 19^th^ century pubarche described by Tanner^18^. The periods for ancestors in the 19^th^ century were set at the ages 8-10 years for female ancestors and at 9-12 years of age for male ancestors. Excess food was not understood to be a problem in the 19^th^ century in Överkalix but to analyze potential transgenerational responses we were obliged to focus on the SGP where we indeed found mismatches reflecting transgenerational responses not only to famine but also to excess food availability. We introduced the concept of the “slow growth period”^1^, the causes and mechanisms of which we discuss here in depth.

### Samples and pedigrees

Eight subjects, seven of which 75 years of age and one 63, consented to blood sampling. They were selected from the 1935 birth cohort in an original epidemiologic study of transgenerational responses to variable availability of food during ancestors’ SGP in Överkalix, Sweden, described in Tinghög et al.^5^ During SGP two paternal grandmothers were exposed to famine (grade 0-2) in 1867 and 1900 respectively, and two to surfeit (grade 5-6) in 1871 and 1879. Two paternal grandfathers were exposed to famine (grade 0-2) in 1867 and 1877 respectively and two were exposed to surfeit (grade 5-6) in 1887 and 1881. These subjects represented four pairs of grandchildren of paternal grandparents exposed to feast or famine (Fig1). Individuals inheriting grandparents who had been exposed to medium crops (grade 3-4) were not included in this strategic sample.

**Figure 1.**
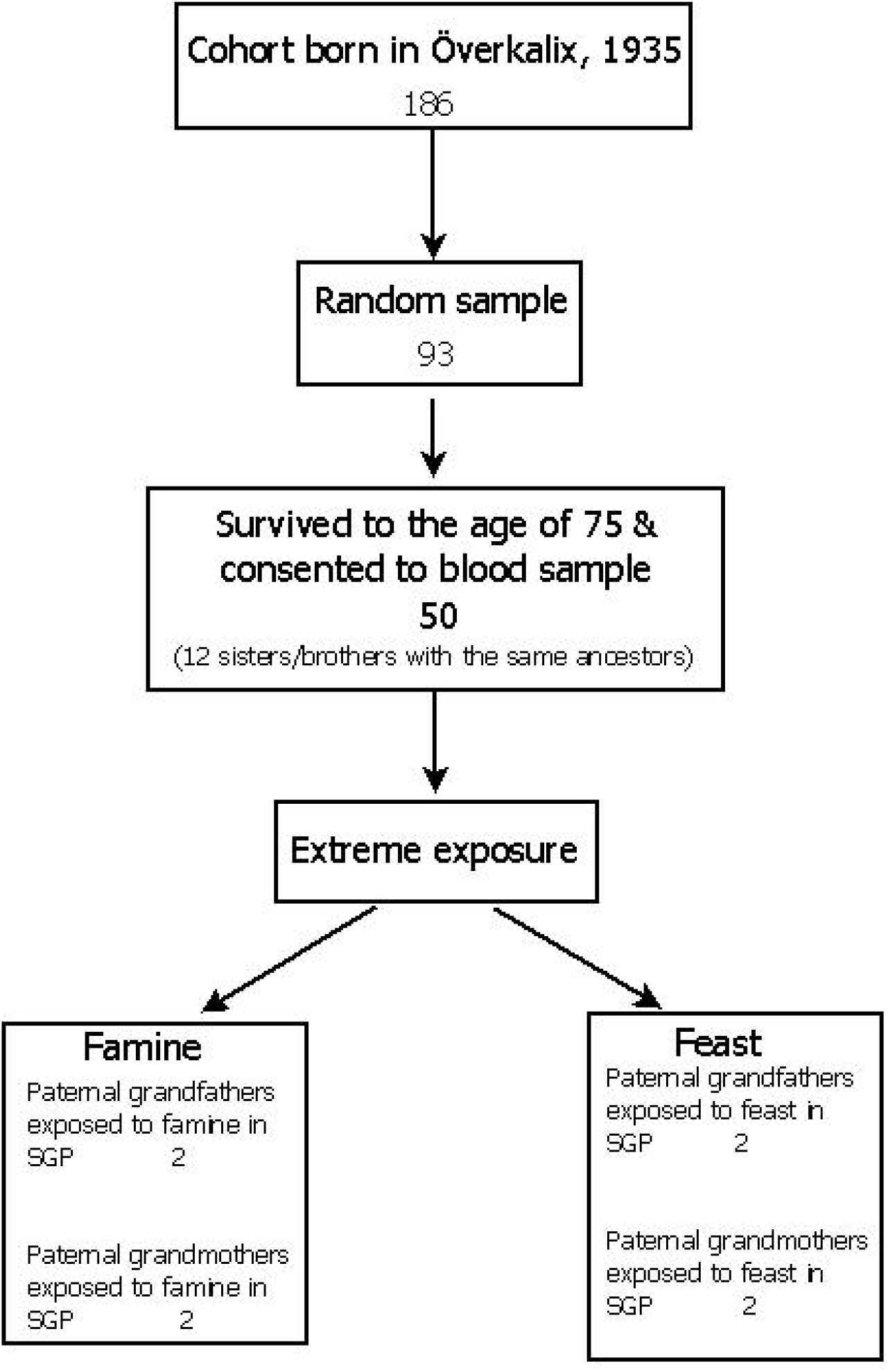
Flowchart

Whole blood samples in EDTA were drawn at county health service units close to the subjects’ residence, frozen at 70° C and sent to the project center. Pedigrees were originally tracked at parish offices in Sweden and Finland and presently in the National archive, the Regional archive of north Sweden, the Research archive of Umeå University, the National tax bureau and digitized genealogical net-based registries. Detailed possible confounders were not identified in this natural experiment: probands, the grandchildren, did not experience any lifetime exposure to famine. The modern food availability though was there from their young adulthood.

### Metabolic pathways posited

The gene ontologies chosen *a priori* described processes related to cardiovascular risks related to hunger, excess food and reactions to environmental stress (http://amigo.geneontology.org). Posited processes during *famine* were “glucose homeostasis” and “glucose transport”, “ketone body biosynthesis and catabolic processes”, “cellular ketone body metabolic pathways”, and “cholesterol homeostasis”. In *hunger and excess* “insulin processing and binding”, “insulin-like growth factor”, “insulin receptor signaling” and “autophagy pathways” were posited. Individuals’ emotional reactions to the stress of famine were assumed to influence hypothalamic and adrenal gland development. Availability and absorption of folate providing methyl donors was of obvious interest, though not the focus of this report. Other posited GOs include appetite, ghrelin, leptin, calcium signaling, eating behavior, adipogenesis, insulin secretion, and lipids.

Altogether, 40 pathways were tested, many of them overlapping and covering three kinds of response pathways reflecting the 3 posited mechanisms.

#### Pathways undergoing unbiased testing

We tested all gene ontology terms in a public database, (UniProt-GOA) including about 40 000 gene ontologies, for grandchildren’s responses to paternal grand parental exposure to poor *versus* excess food.

### Genome-wide methylation

Genomic DNA extraction from whole-blood (buffy coat in one) used the GeneCatcher Blood kit (Invitrogen, Carlsbad, CA, USA). 500 ng of genomic DNA was bisulfite converted with EZ- 96 DNA Methylation kit (Zymo Research, Irvine, CA, USA) and genome wide DNA methylation analysis was carried out using the Infinium MethylationEPIC Bead Chip (IIIumina, San Diego, CA, USA). The Laboratory procedures followed the manufacturers’ protocol.

The array was designed for genome wide methylation analysis with coverage across gene regions with sites in the promoter region, 5UTR, enhancer and gene body, and interrogating more than 850 000 methylation sites at single nucleotide resolution. This technique uses two different probe types (Infinium 1 and 2) with different characteristics, thus requiring normalization to reduce technical bias. GenomeStudio software, version 2011.1 (Illumina Inc.), was used for analysis, visualization and extraction of methylation data.

Methylation levels (beta values) were estimated as the ratio of signal intensity of the methylated alleles to the sum of methylated and unmethylated intensity signals. The beta-values vary from 0 (no methylation) to 1 (100% methylation).

The chip covers CpG islands, shores, shelves and the promoted genes. GOs containing genes and biological pathways were assigned from the literature, primarily from the Gene Ontology project (geneontology.org) chosen for their known intra-generational relation to excess and lack of food.

The assay protocol combined bisulfite conversion of genomic DNA and whole-genome amplification with array-based capture and scoring of the CpG loci. Signal intensity was measured by scanner to generate beta values, the degree of methylation at a locus. Allele-specific single base extension of the probes incorporated a biotin nucleotide ora dinitrophenyl labeled nucleotide. Signal amplification of the incorporated label further improved the overall signal-to-noise ratio of the assay.

Human genomes punctuated by DNA sequences with high frequencies of CpG sites, >200 bases, with CpG content of 50% were termed “*islands*” (CGIs). Differential methylation induced by environment visible in CGIs or in “northern- and southern-*shores* “, within 2 kilo base-pairs (<2kbp) of the islands or in “northern- and southern-*shelves*” within another 2 kilo base pairs from the shores (<2kbp) were studied in CGIs in grandchildren of grandparents exposed to extremes of food availability during their childhood SGP. They are presented as DMRs expressed as percent.

### Statistical analysis

Associations between genes and GO terms were retrieved from the Gene Ontology Association (UniProt-GOA) Database, http://www.ebi.ac.uk/GOA.

Number of genes associated with a GO term and represented with at least one probe on the EPIC chip and number of probes associated with the genes were analyzed among index cases, the grandchildren.

Differentially methylated female index cases whose paternal grandmothers were exposed to bumper harvests in their mid-childhood SGP and to crop failure, respectively, were recorded as well as male index cases whose paternal grandfather had the same extreme exposures.

The criteria for differential methylation was a p-value < 0.05 and an average difference in beta-value between the group averages >0.1. No adjustment for multiple comparisons was carried out. The decreased power of genome-wide analysis, by having negative controls in the remainder of the genome, was compensated by the strict criteria chosen.

Tables were prepared after beta-mixture quantile normalization (BMIQ) correcting probe design bias and with BMIQ-normalized beta values and annotations. The analyses were carried out in Bioconductor-package ChAMP package default probe filtering based on detection p-value, minimum bead count and probes possibly confounded by SNPs whereupon cross hybridization (https://www.ncbi.nim.nih.gov/pubmed/24063430) left 813007 probes for further analysis.

## REFERENCES

1. Bygren LO, Kaati G, Edvinsson E. Longevity determined by paternal ancestors’ nutrition during their slow growth period. Acta Biotheoretica 2001;49:53–59.

2. Kaati G, Bygren LO, Edvinsson S. Cardiovascular and diabetes mortality determined by nutrition during parents’ and grandparents’ slow growth period. Eur J Hum Genet 2002;10:682–8.

3. Pembrey ME, Bygren LO, Kaati G, Edvinsson S, Northstone K et al. Sex-specific, male-line transgenerational responses in humans. Eur J Hum Genet 2006;14:159–166

4. Kaati G, Bygren LO, Pembrey M, Sjostrom M. Transgenerational response to nutrition, early life circumstances and longevity. Eur J Hum Genet 2007; 15:784–90.

5. Tinghög P, Carstensen J, Kaati G, Edvinsson S, Sjöström M, Bygren LO. Migration and mortality trajectories: a study of individuals born in the rural community of Överkalix. Soc Sci Med. 2011;73:744–51.

6. Bygren LO, Tinghög P, Carstensen J, Edvinsson S, Kaati G, Pembrey ME, Sjöström M. Change in paternal grandmothers’ early food supply influenced cardiovascular mortality of the female grandchildren. BMC Genet 2014;15:12.

7. Van den Berg GJ, Pinger PR. Transgenerational effects of childhood conditions on third generation health and educational outcomes. EconHumBiol 2016;23:103–120.

8. Vågerö D, Pinger PR, Aronsson V, van den Berg G. Paternal grandfather’s access to food predicts all-cause and cancer mortality in grandsons. Nat Comm

9. Sales VM, Ferguson-Smith AC, Patti ME. Epigenetic mechanisms of transmission of metabolic disease across generations. Cell Met 2017;25:559–571.

10. Nagy C, Turecki G. Transgenerational epigenetic inheritance: an open discussion. Epigenomics 2015;7:781–790.

11. Hall E, Volkov P, Dayeh T, Esguerra JLS, Salö S et al. Sex difference in the genome-wide DN A methylation pattern and impact on gene expression, microRNA levels and insulin secretion in human pancreatic islets. Genome Biology 2014;15:522–544.

12. Guo YT, Luo HV, Wu YM, Magdalou J, Chen LB, Wang H. Influencing factors, underlying mechanisms and interactions affecting hypercholesterolemia in adult offspring with caffeine exposure during pregnancy, Reprod Toxicol 2018;79:47–56

13. Earon SA, Jayasooriah N, Buckland ME, Buckland ME, Martin DI, Cropley JE, Suter CM. Roll over Weismann: extracellular vesicles in the transgenerational transmisson of environmental effects Epigenomics 2015; 7:1165–71.

14. Vaiserman AM, Koliada AK, Jirtles RL. Non-genomic transmission of longevity between generations: potential mechanisms and evidence across species. Epigenetics Chromatin 2017;D0I 10.1186/s 13072-017-0145-1

15. Hellstenius J. Skördarna i Sverige och deras verkningar. [Harvests in Sweden and their repercussions]. Statistisk Tidskrift 77–119. Stockholm, Sweden 1871 (sic).

16. Jörberg L. (1972). A history of prices in Sweden 1732-1914. CWK Gleerup, Lund, Sweden.

17. Tanner J.M. A History of the Study of Human Growth. Cambridge University Press, Cambridge 1981

18. Prader, A., R.H. Largo, L. Molinari and C. Issler (1989). Physical growth of Swiss children from birth to 20 years of age. First Zurich Longitudinal Study of Growth and Development. Helvetica Paediatrica Acta 43:Suppl 52: 1–125

